# Microtubule detyrosination enhances matrix remodeling and force transmission during endothelial sprouting

**DOI:** 10.64898/2026.01.06.698015

**Authors:** Danahe Mohammed, Ibrahim Hamid, Sarah Steffens, Charlotte Lemeslier, Valerie Martinelli, Apeksha Shapeti, Hans Van Oosterwyck, Benoit Vanhollebeke, Sylvain Gabriele, Maud Martin

## Abstract

Angiogenic sprouting relies on the coordinated regulation of endothelial migration, force generation, and extracellular matrix (ECM) remodeling. While microtubules are known to regulate cell polarity and migration, how tubulin post-translational modifications reshape microtubule behavior and impact matrix-dependent migration programs remain unclear. Here, using sprouting angiogenesis as an integrative physiological context, we combine live-cell imaging, three-dimensional (3D) sprouting and microfluidic invasion assays, and 3D traction force microscopy to dissect the role of tubulin detyrosination and acetylation. We show that microtubule detyrosination, in contrast to acetylation, promotes directional persistence, sprout elongation and collective endothelial invasion. Detyrosinated microtubules remain dynamic and display more persistent growth, supporting sustained directional migration. Strikingly, microtubule detyrosination also enhances ECM remodeling, leading to increased matrix degradation and augmented traction force transmission. Functional inhibition experiments reveal that matrix degradation, rather than actomyosin contractility, is required for both elevated forces and enhanced sprouting. Together, these findings identify microtubule detyrosination as a key regulator linking cytoskeletal dynamics, cell migration and matrix remodeling during angiogenesis.

## Introduction

Angiogenic sprouting is a fundamental process in vascular morphogenesis, requiring the tight coordination of endothelial cell migration, force generation, and extracellular matrix remodeling^1–3^. This highly dynamic behavior depends on the cytoskeleton, which integrates biochemical and mechanical cues to drive collective invasion and lumen formation in three dimensional (3D) environments. While the role of actin filaments in angiogenesis^4–6^ has been extensively studied, the contribution of microtubules has only recently begun to emerge^7^. Microtubules not only serve as structural tracks for vesicle trafficking^8^ and organelle positioning^9^ but also act as critical regulators of force transmission and mechanosensitive signaling during migration^10–12^, including in endothelial responses to mechanical stimuli^13,14^. Notably, non-centrosomal microtubules have been shown to be essential for endothelial polarization during sprouting angiogenesis^7^.

A growing body of work indicates that tubulin post-translational modifications (PTMs) confer functional specialization to microtubules^15^. Distinct PTMs, including detyrosination and acetylation, form part of a so-called "tubulin code" that modulates microtubule stability, motor protein processivity, and the recruitment of specialized microtubule-associated proteins (MAPs)^15^. Detyrosinated microtubules are typically associated with long-lived, mechanically resilient filaments^16^, and preferentially recruit kinesin-1^17–19^ and kinesin-2 family motors^20,21^. In contrast, MAPs containing CAP-Gly domains, such as the dynactin subunit p150Glued^22^ and CLIP170^23^, selectively recognize the tyrosine-containing C-terminal motif of α-tubulin, while depolymerizing kinesins MCAK and KIF2A display higher activity on tyrosinated lattices^24^. Acetylation, which occurs within the microtubule lumen, is likewise enriched on stable microtubules^25^ and enhances resistance to mechanical breakage under compressive stress^26^.

Recent studies have begun to link these PTMs to cell polarity, adhesion dynamics and mechanosensitivity, processes central to endothelial behavior during angiogenesis. Detyrosinated microtubules have been shown to support cell polarization in epithelial systems^27^, whereas microtubule acetylation enhances focal adhesion dynamics and modulates mechanosensitivity responses in astrocytes and endothelial cells^11^. Moreover, microtubule acetylation alone has been reported to promote collective cell migration in the neural crest^28^. In the vascular context, acetylated microtubules are required for endothelial elongation and alignment under shear stress ^14^, while reduced microtubule detyrosination leads to vascular patterning defects in the zebrafish trunk^29^. Despite these advances, how distinct tubulin PTMs differentially regulate endothelial sprouting and matrix-dependent invasion remains largely unexplored.

Here, we address this knowledge gap by generating cellular models in which microtubules are uniformly modified, enabling controlled interrogation of the respective roles of tubulin detyrosination and acetylation in endothelial behavior. Using complementary 3D sprouting and microfluidic invasion assays combined with live-cell imaging and 3D traction force microscopy, we uncover a striking functional divergence between these two modifications. While both are classically associated with stable microtubules, detyrosination strongly promotes endothelial sprouting, whereas acetylation is detrimental. Detyrosinated microtubules remain dynamic and display persistent growth, supporting sustained directional migration. They also promote matrix metalloproteinase (MMP)-dependent extracellular matrix degradation that enhances force transmission, collectively sustaining efficient multicellular sprouting.

This work identifies tubulin detyrosination as a central cytoskeletal regulator coordinating microtubule dynamics, force transmission and matrix remodeling during endothelial invasion, thereby providing a mechanistic framework for vascular sprouting in regenerative and pathological contexts.

### Microtubule detyrosination enhances invasion and traction forces during multicellular angiogenic sprouting

To dissect the contribution of tubulin PTMs to angiogenic sprouting, we engineered cellular models with distinct and uniform microtubule modification states using human umbilical vein endothelial cells (HUVECs) **(Fig.1A-D)**. Control cells (CTRL) displayed negligible levels of microtubule detyrosination (0.44% ± 0.36) and modest acetylation (23.63% ± 6.31). To selectively enhance these reversible PTMs, we shifted the balance towards modification by forcing the stable expression of the corresponding modifying enzymes **(Fig. 1A)**. Overexpression of the VASH1-SVBP tubulin carboxypeptidase complex^30,31^ generated a *HyperDetyr* model characterized by a near-complete detyrosination of the microtubule network (97.72% ± 1.56), without affecting acetylation levels (22.57% ± 9.98) **(Fig. 1C,D)**. Conversely, overexpression of the acetyltransferase αTAT1^32,33^ yielded a *HyperAcetyl* model, marked by a strong increase in microtubule acetylation (79.58% ± 17.29), while maintaining minimal detyrosination (0.25% ± 0.22) **(Fig. 1C,D)**.

**Fig. 1.**
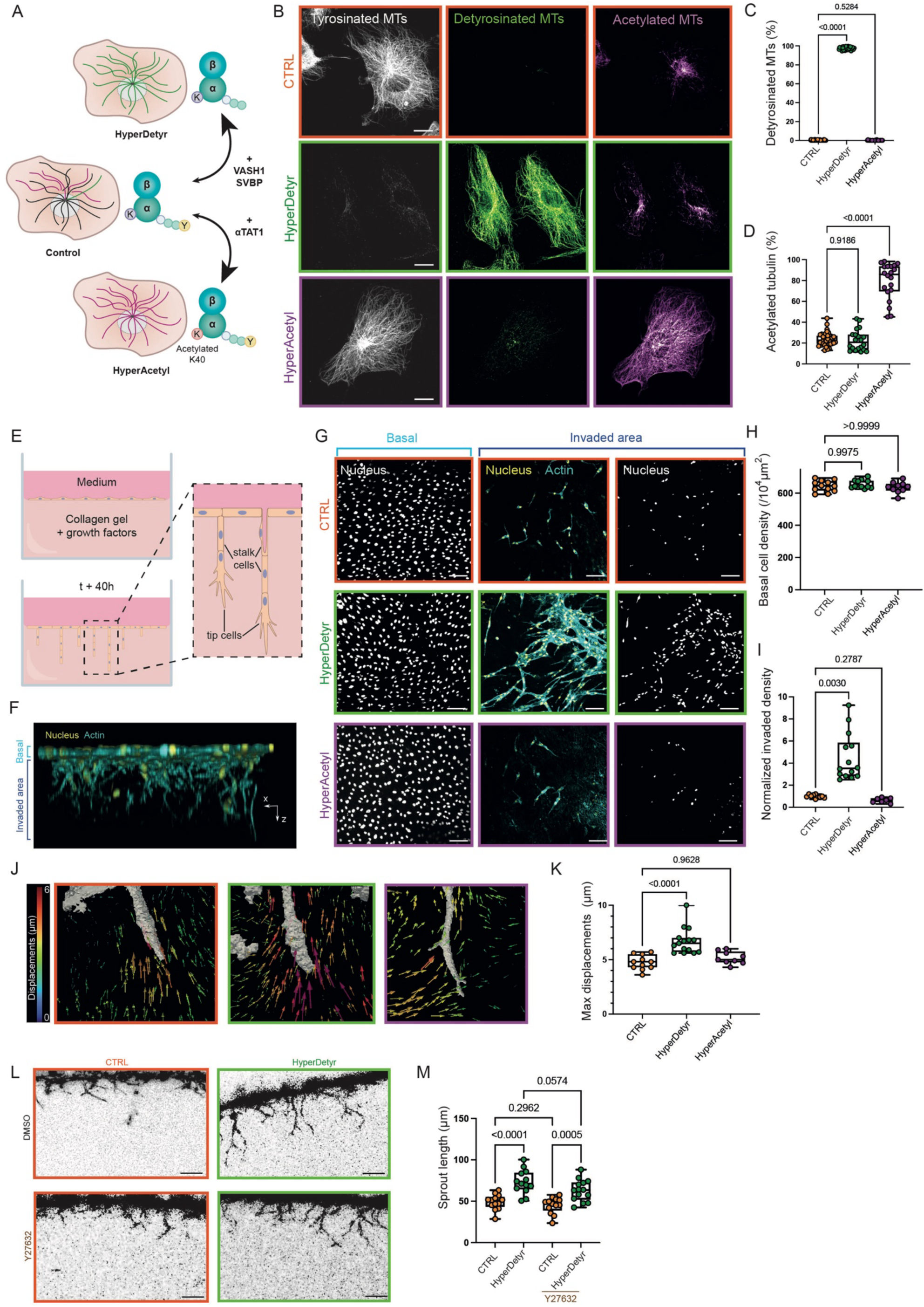
Microtubule detyrosination enhances invasion and traction forces during multicellular angiogenic sprouting. (A) Schematic representation of the strategy used to modulate PTMs. Overexpression of ATAT1 was used to induce microtubule hyperacetylation, whereas overexpression of the VASH1-SVBP complex was employed to promote microtubule hyperdetyrosination in HUVECs. (B) Representative immunofluorescence images of HUVECs displaying differential enrichment of microtubule acetylation and detyrosination. Three cellular models are generated with modulation of PTMs: control cells (CTRL), *HyperDetyr* and *HyperAcetyl* cellular models. Cells were stained for tyrosinated (gray), detyrosinated (green), and acetylated (purple) tubulin. Scale bar=5 µm. (C) The control (CTRL, orange) and *HyperAcetyl* (purple) cells display negligible levels of detyrosinated microtubules, whereas the *HyperDetyr* model (green) exhibits near-complete microtubule detyrosination. The signal intensity of the detyrosinated tubulin signal was quantified and normalized to the total tubulin signal (n>30, N>6 independent experiments). (D) The CTRL and *HyperDetyr* cells show approximately 20% acetylated microtubules, while the *HyperAcetyl* cells exhibit near-complete microtubule acetylation. The signal intensity of the acetylated tubulin signal was quantified and normalized to the total tubulin signal (n>20, N>6 independent experiments). (E) Schematic representation of the 3D endothelial sprouting assay. (F) Confocal 3D reconstruction of a sprouting assay of CTRL model showing the basal monolayer (top) and the invaded sprouting (bottom). Nuclei were stained with DAPI (yellow) and sprouts with Phalloidin (labeling F-actin, cyan). (G) Representative maximum intensity z-projection of nuclei in the basal area (light blue) and invaded area (dark blue) for CTRL, *HyperDetyr* and *HyperAcetyl* cells. Stainings as in F. Scale bar=100 µm. (H) Basal cell density remains unchanged across conditions. The number of nuclei per 10^4^ µm^2^ was counted in the monolayer (N>10 independent experiments). (I) Invaded cell density is significantly increased in the *HyperDetyr* condition compared to CTRL and *HyperAcetyl* models. The number of nuclei per 10^4^ µm^2^ was counted in the invaded area (N>7 independent experiments). (J) Representative 3D displacement maps computed around endothelial sprouts after 18 h of invasion. (K) Maximum displacement values are higher in the *HyperDetyr* model compared to CTRL. The bead displacements were quantified (see Methods for details) and the maximum value per sprout was selected (N>7 independent experiments). (L) Representative maximum intensity projections of angiogenic sprouts stained with phalloidin after 6 h of invasion in CTRL and *HyperDetyr* models treated with DMSO or Y27632 10µM (a ROCK inhibitor). Scale bar=50 µm. (M) Treatment with Y27632 does not affect sprout length in any of the three cellular models. The sprout average length was quantified in each condition (n>13, N>6 independent experiments).

To quantitatively assess the extent of these modifications beyond bulk intensity measurements, we measured the fraction of the microtubule lattice covered by each PTM. Detyrosination spanned 98.59% ± 0.16 of the lattice in *HyperDetyr* cells, while acetylation covered 90.44% ± 6.72 of the lattice in *HyperAcetyl* cells **(Extended Fig. 1A, B)**. Together, these models act as molecular gatekeepers that selectively turn one microtubule modification fully “on” while keeping the other “off”, enabling functional interrogation of detyrosination and acetylation in a controlled cellular context.

We next applied this experimental framework to assess how distinct microtubule PTMs influence endothelial invasion using a 3D endothelial sprouting assay **(Fig. 1E-I)**. HUVECs were seeded as a confluent monolayer facing a growth factor-rich soft collagen matrix, allowing sprouts to invade the extracellular matrix in a configuration that recapitulates early vessel formation. Confocal imaging and 3D reconstruction enabled separate segmentation of the basal monolayer and invading sprouts (**Fig. 1F)**. While basal cell density remained comparable across all conditions **(Fig. 1G, H)**, *HyperDetyr* cells exhibited a striking ∼4.5-fold increase in invading cell density compared with CTRL and *HyperAcetyl* cells (**Fig. 1G, I, and Extended Fig. 1C-E)**. Importantly, these resulting sprouts were multicellular and formed well-defined lumens **(Extended Fig. 1C, F)**, indicating that detyrosination enhances invasive sprouting without disrupting collective organization. In contrast, *HyperAcetyl* cells displayed a modest but significant reduction in invasion relative to CTRL, revealing an effect opposite to that of detyrosination **(Fig. 1G, I, and Extended Fig. 1E)**.

The pro-invasive effect of microtubule detyrosination was further supported by competitive co-culture experiments **(Extended Fig. 1G)**. When CTRL and *HyperDetyr* cells were mixed at equal ratios, invasion increased compared with CTRL alone without reaching the level of pure *HyperDetyr* cultures **(Extended Fig. 1G, H)**. Strikingly, nearly all invading cells (88.83% ± 7.52) correspond to *HyperDetyr* cells **(Extended Fig. 1G, I)**, underscoring a strong benefit conferred by microtubule detyrosination. Time-lapse imaging revealed that this advantage amplified over time: differences were modest at early stages (∼1.2-fold at 8 h) but became increasingly pronounced at later time points (∼1.9-fold at 24 h ∼2.1-fold at 40 h; **Extended Fig. 1J, K**).

The magnitude of the enhanced invasion observed in *HyperDetyr* cells prompted us to ask whether increased traction forces might explain their hyperinvasive behavior^2,34^. We therefore quantified sprout-generated forces using 3D traction force microscopy (TFM)^34^ **(Fig. 1J, K and Extended Fig. 1L, M)**. Endothelial sprouts enriched in detyrosinated microtubules exerted significantly larger matrix deformations, with increased maximal displacements (6.70 µm ±1.20) compared to CTRL (4.75 µm ±0.67), whereas *HyperAcetyl* cells showed no significant changes (5.16 µm ±0.58) **(Fig. 1J, K)**. These results indicate that microtubule detyrosination enhances force generation during sprouting angiogenesis.

Because actomyosin contractility is classically considered as the primary driver of cellular traction forces, we tested its contribution to the detyrosination-dependent phenotype by inhibiting ROCK-mediated contractility with Y-27632^35^. Surprisingly, ROCK inhibition did not reduce sprout length or invasion in *HyperDetyr* cells (**Fig. 1L, M**), indicating that the enhanced sprouting and force transmission associated with microtubule detyrosination occur independently of actomyosin contractility. Together, these findings suggest that detyrosinated microtubules promote angiogenic invasion through a distinct mechanism, pointing to alternative modes of force transmission that we investigate next.

### Microtubule detyrosination promotes migratory persistence independently of tip cell selection and proliferation

Analysis of sprouting kinetics revealed that early invasion dynamics were comparable across microtubule PTM conditions **(Extended Fig. 1J, K)**, indicating that tubulin modifications do not affect sprout initiation. To confirm this, we used an orthogonal side-view sprouting assay to quantify both sprout number and cellular content at early (8 h) and later (24 h) time points **(Extended Fig. 2A-C)**. Sprout numbers were almost unchanged between CTRL and *HyperDetyr* conditions at both time points **(Extended Fig. 2A, B)**. In contrast, *HyperDetyr* sprouts contained significantly more cells at t=24 h (≈ 1.5-fold relative to CTRL, **Extended Fig. 2A, C**) and were consistently longer at both 8 h and 24 h **(Extended Fig. 2A, D)**. Thus, the enhanced invasion observed upon microtubule detyrosination arises from longer multicellular sprouts rather than generation of additional sprouts.

During physiological angiogenesis, the selection of tip cells for sprouting is tightly controlled by vascular endothelial growth factor (VEGF) signaling, with NOTCH signaling further restricting neighboring stalk cells from adopting the tip fate. Inhibition of NOTCH signaling leads to excessive tip cells and a hypersprouting phenotype^36,37^. To determine whether microtubule detyrosination affects this process, we blocked NOTCH activity with the γ-secretase inhibitor DAPT^38^ during sprouting **(Fig. 2A-C)**. DAPT treatment increased sprout number in both CTRL and *HyperDetyr* conditions, while sprout length remained unchanged **(Fig. 2A-C)**, indicating that detyrosinated cells retain sensitivity to canonical angiogenic signaling cues. Importantly, the invasive advantage of *HyperDetyr* cells persisted under condition of relieved lateral inhibition **(Fig. 2A, C)**. Consistently, 1:1 co-culture assays further demonstrated that detyrosination does not bias tip-cell selection: after 6 h of sprouting, tip cells were equally distributed between CTRL and *HyperDetyr* populations **(Extended Fig. 2E, F)**. We next asked whether increased proliferation could account for the larger size of *HyperDetyr* sprouts. However, proliferation rates were comparable across conditions (**Fig. 2D)**, excluding cell division as a major contributor. We therefore turned to cell motility, a central determinant of angiogenic sprouting. Collective migration was first assessed using a 2D scratch assay. While wound closure was similar between CTRL and *HyperDetyr* cells, migration was significantly impaired in *HyperAcetyl* cells **(Extended Fig. 2G,H)**, consistent with previous reports showing that elevated microtubule acetylation hampers efficient cell migration^11^ and invasion^37^.

**Fig. 2.**
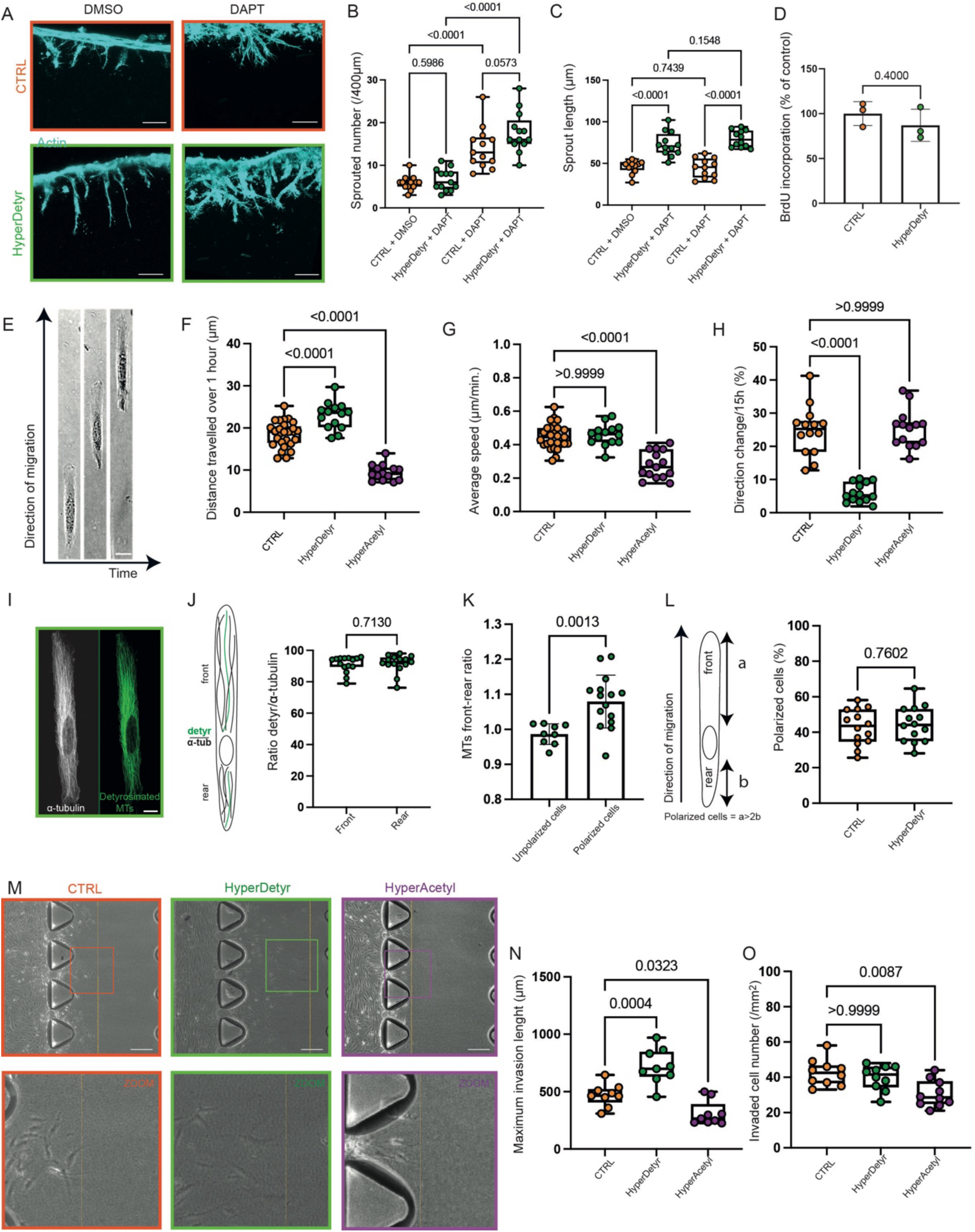
Microtubule detyrosination promotes migratory persistence. (A) Representative confocal images showing hypersprouting in the presence of DAPT (NOTCH inhibitor) (right). Sprouts were stained with Phalloidin (labeling F-actin, cyan). Scale bar=50 µm. (B) Both CTRL and *HyperDetyr* models exhibit increased sprout number upon DAPT treatment (N=13 independent experiments). (C) Sprout length is unaffected by DAPT treatment in both conditions and remains consistently longer in the *HyperDetyr* model (N=13 independent experiments). (D) Proliferation assay comparing CTRL and *HyperDetyr* models. The proliferation rate does not differ between the two conditions (N=3 independent experiments). (E) Brightfield images of HUVECs migrating along microprinted lines. (F) Quantification of the distance traveled over 1 hour (N=14 independent experiments). (G) Quantification of average migration speed shows no difference between CTRL, *HyperDetyr* and *HyperAcetyl* models (N=14 independent experiments). (H) *HyperDetyr* cells, however, exhibit increased directional persistence (N=14 independent experiments). (I) Immunofluorescence images of α-tubulin (white) and detyrosinated microtubules (green) in *HyperDetyr* cells migrating on micropatterned lines. Scale bar=2 µm. (J) Quantification of the ratio of detyrosinated microtubules in the front and rear of the cells shows no anterior enrichment (N=15 independent experiments). (K) In contrast, total microtubules (α-tubulin) are enriched at the front of the cells. (L) The proportion of polarized cells does not differ between CTRL and *HyperDetyr* models (N>9 independent experiments). A cell was considered polarized if the front (a) was at least twice the rear axis (b). (M) Brightfield image of the invasion assay performed under VEGF gradient conditions. The dashed yellow line indicates the maximal invasion front. Scale bar = 200 µm. (N) The *HyperDetyr* model exhibits a greater invasion distance compared to CTRL (N=9 independent experiments). (O) Despite the increased invasion length, the total number of invading cells is comparable between CTRL and *HyperDetyr* conditions. The maximum invasion length and invaded cell number were quantified in each condition (N=9 independent experiments).

To more closely mimic the spatially constrained environment of 3D migration, we next analyzed cell behavior on 1D micropatterned lines, a system that recapitulates core features of 3D migration^27,39^, and performed live tracking of cells **(Fig. 2E)**. In agreement with the sprouting assays, *HyperDetyr* cells increased the distance travelled by cells over a one-hour period, whereas microtubule acetylation reduced it **(Fig. 2F)**. Detailed analysis of migration parameters revealed distinct underlying mechanisms: *HyperDetyr* cells displayed a pronounced increase in directional persistence, characterized by a marked reduction in direction changes (6.06% ± 2.89) compared with CTRL cells (24.71% ± 7.45), whereas *HyperAcetyl* cells exhibited reduced migration speed **(Fig. 2G, H)**. These results indicate that microtubule detyrosination enhances migration persistence rather than migration velocity.

To investigate the basis of this increased migration persistence, we examined microtubule organization in polarized cells migrating on micropatterned 1D lines. Previous studies reported that enrichment of detyrosinated microtubules at the leading-edge supports directed migration^27^. However, as expected from a uniformly detyrosinated microtubule network **(Extended Fig.1B)**, *HyperDetyr* cells did not display front-rear asymmetry in detyrosinated microtubules **(Fig. 2I, J)**. Instead, total microtubules were significantly enriched at the leading edge **(Fig. 2K)**, suggesting that global microtubule enrichment, rather than spatially biased detyrosination, is sufficient to sustain polarity. Consistently, the proportion of polarized cells was comparable between CTRL and *HyperDetyr* populations **(Fig. 2L)**, indicating that detyrosination does not enhance polarization but stabilizes directional migration once polarity is established.

*In vivo*, endothelial cells migrate directionally in response to chemotactic cues such as VEGF. To test whether tubulin PTMs modulate chemotactic migration, we performed 3D chemotaxis assays using a microfluidic device that generates a stable VEGF gradient across a collagen matrix^40^. This setup induced robust cell invasion toward the VEGF source **(Extended Fig. 2I-L)**. *HyperDetyr* cells exhibited a marked increase in maximal invasion length compared with CTRL cells **(Fig. 2M, N)**, without a change in the number of invading cells **(Fig. 2O)**. In contrast, *HyperAcetyl* cells showed fewer invading cells that migrated shorter distances **(Fig. 2M-O)**. Morphological analysis revealed that *HyperDetyr* cells formed fewer but longer protrusions at both early and late time points **(Extended Fig. 2M-P)**, consistent with a migratory strategy favoring elongated, persistent protrusions.

Collectively, these results demonstrate that microtubule detyrosination enhances endothelial invasion by promoting migratory persistence independently of tip-cell selection and proliferation. By stabilizing directional migration and polarized protrusive activity, detyrosinated microtubules provide a mechanistic link between cytoskeletal organization and sustained multicellular sprouting.

### Detyrosination preserves microtubule dynamics while promoting persistent growth

The increased migratory persistence observed in *HyperDetyr* cells prompted us to examine whether detyrosination alters the intrinsic properties of microtubules^41^. Tubulin PTMs have long been correlated with microtubule longevity and are often assumed to stabilize the microtubule lattice^42^. However, disentangling intrinsic effects of PTMs from secondary consequences of mixed modification states has remained challenging. Our cellular models, in which microtubules are uniformly detyrosinated or acetylated, provide a unique framework to directly address this question.

We first assessed microtubule stability using complementary depolymerization assays, including nocodazole- and cold-induced depolymerization, as well as the StableMARK live sensor^43^, which reports lattice expansion states associated with microtubule stability in intact cells^44^. Acetylated microtubules were consistently less stable than CTRL across all assays **(Extended Fig. 3A-F)**. In contrast, detyrosinated microtubules displayed context-dependent behavior: they were modestly more resistant to nocodazole-induced depolymerization yet more sensitive to cold treatment **(Extended Fig. 3A-D)**. Consistently, StableMARK labeling revealed fewer stabilized microtubules in *HyperDetyr* cells **(Extended Fig. 3E-F)**. Together with previous reports^43,45,46^, these results indicate that tubulin detyrosination does not confer intrinsic lattice stabilization and that PTMs alone are insufficient to predict global microtubule stability.

The increased resistance of detyrosinated microtubules to nocodazole suggested that detyrosination might preferentially affect plus-ends behavior rather than lattice stability. To test this possibility, we examined microtubule growth dynamics by visualizing EB1 comets, which label actively growing microtubule plus-ends^47^. The density of EB1 comets was unchanged in *HyperDetyr* cells compared with CTRL, indicating preserved microtubule dynamics, whereas comet density was reduced in *HyperAcetyl* cells **(Fig. 3A)**. Because a small fraction of tyrosinated tubulin may persist in *HyperDetyr* cells, we next assessed whether EB1 comets preferentially associated with residual tyrosinated segments. Spatial analysis revealed that ∼96% of comets colocalized with detyrosinated microtubules **(Fig. 3B, C)**, demonstrating that active growth occurs independently of tyrosinated tubulin tips.

**Fig. 3.**
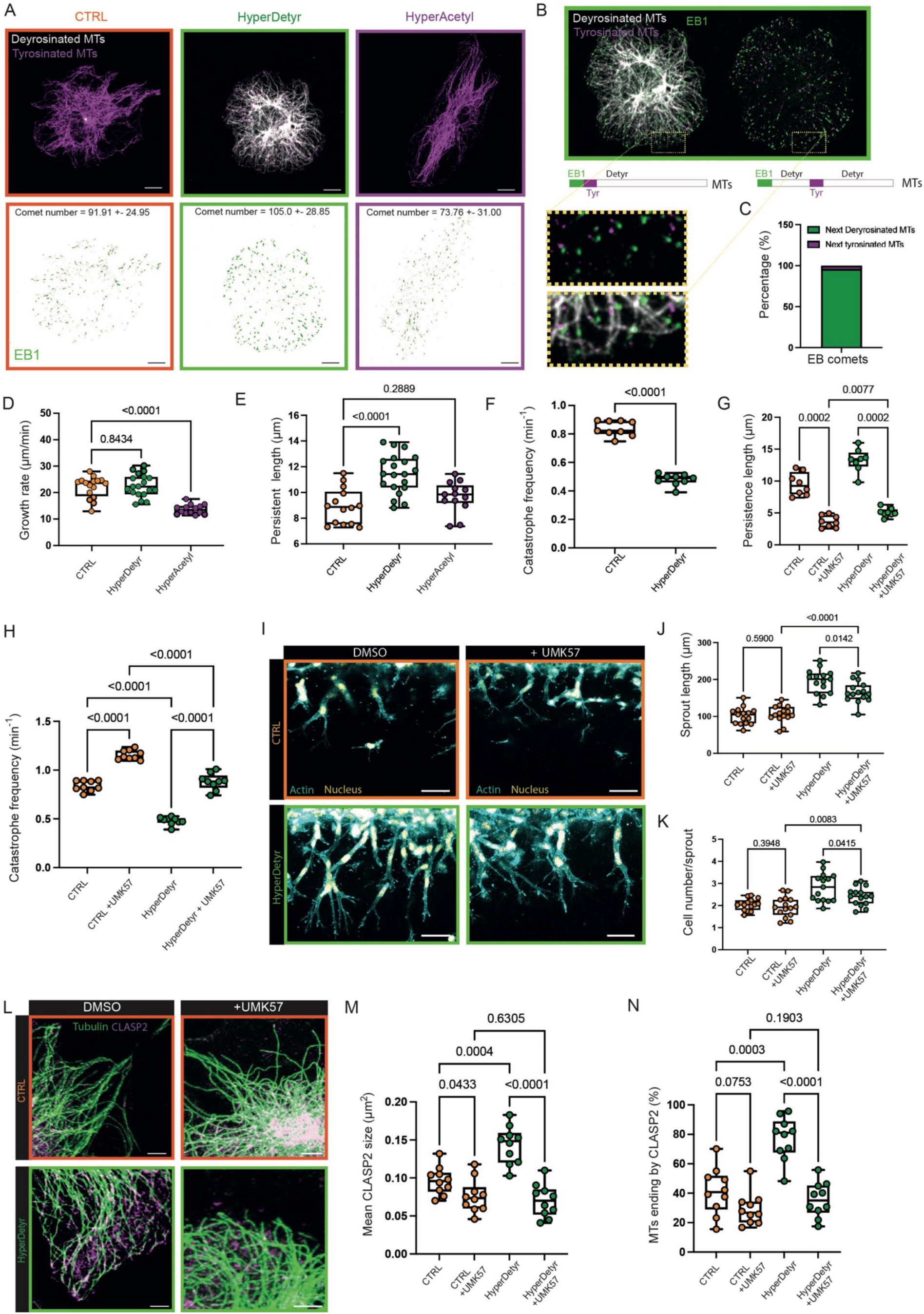
Detyrosination preserves microtubule dynamics while promoting persistent growth. (A) Representative confocal images showing EB1 comets, tyrosinated microtubules and detyrosinated microtubules in CTRL, *HyperDetyr* and *HyperAcetyl* models (scale bar=5 µm) and (B) a Zoom-in view of a *HyperDetyr* model. p value of comet number for CTRL vs HyperDetyr is 0,2038 and 0,0349 for CTRL vs *HyperAcetyl* cells. Cells were stained for detyrosinated (gray), tyrosinated (magenta) tubulin and EB1 (green). Dashed yellow rectangles indicate the areas selected for higher-magnification views shown above. (C) The percentage of EB1 comets associated with tyrosinated or detyrosinated microtubules was measured. In *HyperDetyr* cells, approximately 90% of EB1 comets localize near detyrosinated microtubules. (D) Microtubule growth rate remains unchanged between CTRL and *HyperDetyr* models but is reduced in the *HyperAcetyl* condition (n=16, N>3 independent experiments). (E) The persistence length of microtubules from EB3-mCherry polymerization initiation to termination is increased in the *HyperDetyr* model but unchanged in the *HyperAcetyl* model (n>13, N>3 independent experiments). (F) Catastrophe frequency is strongly decreased in *HyperDetyr* model (n=9, N=3 independent experiments). (G) Persistence length of microtubules for CTRL and *HyperDetyr* model in presence of DMSO or UMK57, an MCAK activator (n=9, N=3 independent experiments). (H) Catastrophe frequency from (F) + catastrophe frequency in presence of UMK57. Catastrophe frequency increases upon addition of UMK57, in both CTRL and *HyperDetyr* models (n=9, N=3 independent experiments). (I) Representative confocal images of sprouting assays, shown in lateral view, in the presence of DMSO (left) or UMK57 (right) for CTRL and *HyperDetyr* models. Nuclei were stained with DAPI (yellow) and sprouts with Phalloidin (labeling F-actin, cyan). Scale bar=50 µm. (J, K) Sprout length (J) and the number of cells per sprout (K) decrease upon UMK57 treatment in the *HyperDetyr* model, whereas no change is observed in CTRL (n=15, N=5 independent experiments). (L) Representative confocal images of microtubules (green) and CLASP2 (purple) in the presence of DMSO or UMK57 for CTRL and *HyperDetyr* models. Cells were stained for α-tubulin (green) and CLASP2 (magenta). Scale bar=3 µm. (M, N) Mean CLASP2 puncta size (M) and the percentage of microtubule ending at CLASP2 (N) both decrease upon UMK57 treatment in CTRL and *HyperDetyr* models (n=10, N=3 independent experiments).

Live-cell imaging of EB1 dynamics further revealed that microtubule growth rates were unchanged in *HyperDetyr* cells but significantly reduced in *HyperAcetyl* cells **(Fig. 3D)**, consistent with reduced dynamicity in the latter condition. Strikingly, detyrosination selectively increased growth persistence, whereas acetylation had no effect **(Fig. 3E)**. This increased persistence could be explained by a reduced catastrophe frequency in *HyperDetyr* cells **(Fig. 3F)**. Thus, detyrosinated microtubules remain dynamic and sustain growth over longer distances, a property well suited to support persistent directional migration.

One known consequence of microtubule detyrosination is a reduced susceptibility to depolymerization by the kinesin-13 family member MCAK^24^. Consistent with the idea that reduced MCAK activity contributes to the increased growth persistence of detyrosinated microtubules, treatment with the MCAK activator UMK57 was able to restore normal microtubule dynamics towards CTRL levels, reducing growth persistence and normalizing catastrophe frequency **(Fig. 3G, H)**. Notably, restoring normal microtubule dynamics partially suppressed the enhanced sprouting capacity of *HyperDetyr* cells **(Fig. 3I-K)**, indicating that persistent microtubule growth is an important factor driving this phenotype, although it likely acts in concert with additional cellular mechanisms.

In addition to MCAK, CLASP proteins are known regulators of microtubule plus-ends stability and growth persistence^48,49^. CLASP2 is of particular interest as it has recently been shown to preferentially associate with detyrosinated microtubules within the mitotic spindle^50^ and has emerged as a critical player in migration, particularly in confined 3D environments, by locally tuning microtubule behavior^12,51,52^. Quantitative confocal image analysis revealed a pronounced increase in CLASP2 association with the detyrosinated microtubule network **(Extended Fig. 3G-I)**. *HyperDetyr* cells displayed both a higher number of CLASP2-positive microtubule ends and larger CLASP2 accumulations, indicating enhanced recruitment of CLASP2 to plus-ends **(Extended Fig. 3G-I)**. These effects were specific to the *HyperDetyr* condition, as they were not detected in the *HyperAcetyl* model, and they did not extend to the Golgi-associated pool of CLASP2, which participates in the formation of Golgi-derived microtubules **(Extended Fig. 3J-L)**.

To determine whether CLASP2 accumulation at plus-ends was directly driven by detyrosination or instead reflected altered microtubule dynamics, we reduced growth persistence in *HyperDetyr* cells using UMK57. Under these conditions, CLASP2 enrichment at microtubule plus-ends was lost **(Fig. 3L-N)**, indicating that CLASP2 accumulation depends on the persistent growth behavior conferred by detyrosination rather than on detyrosination alone. Together, both diminished MCAK activity and increased CLASP2 recruitment likely cooperate to generate a microtubule array characterized by sustained growth persistence in *HyperDetyr* cells.

Collectively, these findings demonstrate that microtubule detyrosination preserves dynamic instability while selectively promoting persistent microtubule growth. This emergent dynamic state provides a mechanistic basis for the enhanced migratory persistence observed in detyrosinated endothelial cells.

### Matrix degradation governs enhanced sprouting in *HyperDetyr* cells

Normalizing microtubule dynamics only partially corrected the *HyperDetyr* sprouting phenotype, indicating that additional mechanisms contribute to enhanced angiogenic sprouting. Because endothelial invasion critically depends on cell-matrix interactions, we next examined whether microtubule detyrosination modulates extracellular matrix (ECM) remodeling.

To directly assess matrix degradation activity, we performed fluorescent gelatin degradation assays. *HyperDetyr* cells exhibited a markedly increased degraded area (1.35 ± 0.63) compared with CTRL cells (0.58 ± 0.09), whereas *HyperAcetyl* cells showed no increase (0.68 ± 0.25) **(Fig. 4A, B)**. Notably, degradation activity positively correlated with the magnitude of traction forces exerted on the matrix, as quantified by maximal displacements in 3D TFM **(Fig. 1J, K)**, suggesting a functional coupling between ECM remodeling and force transmission **(Fig. 4C)**.

**Fig. 4.**
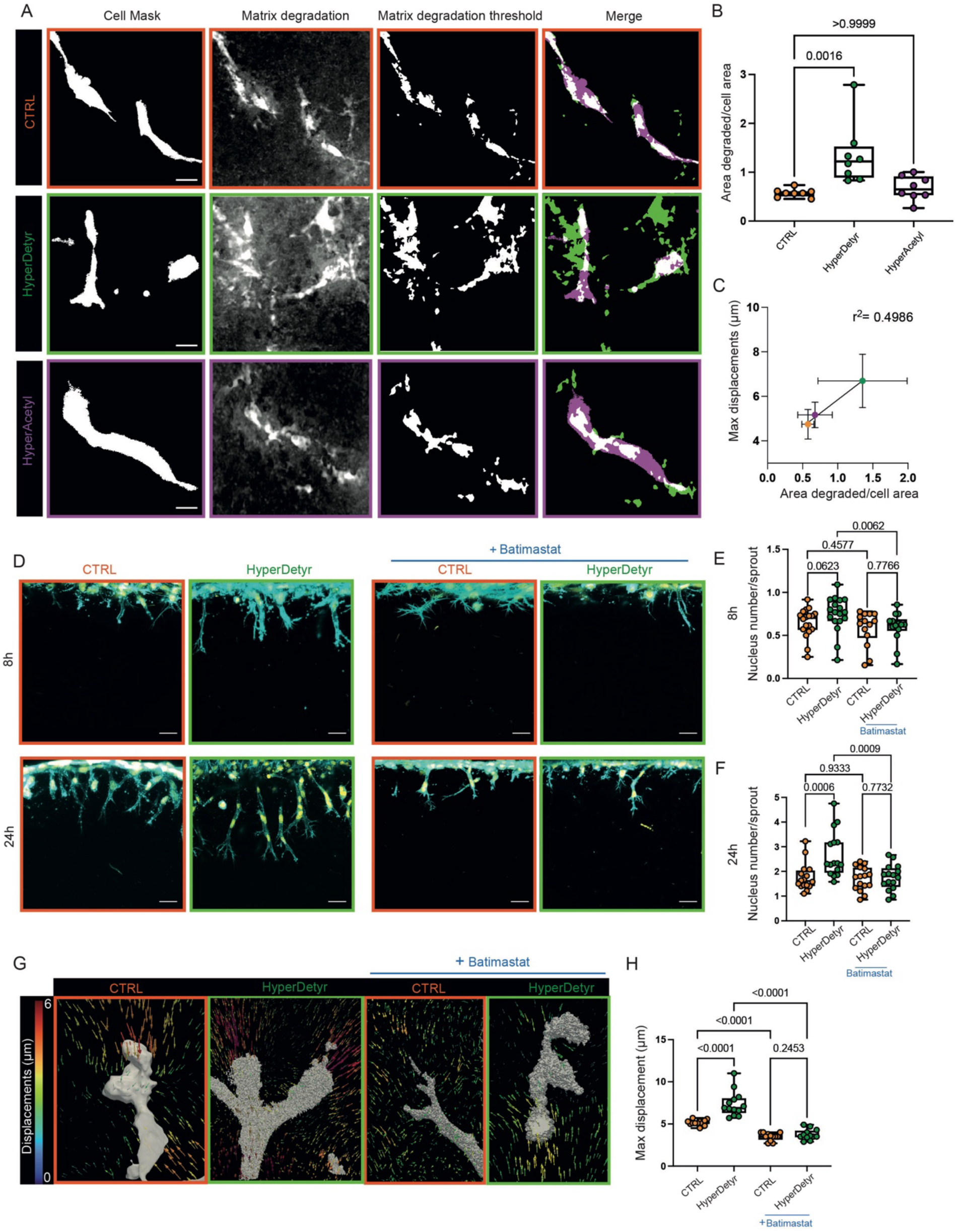
Matrix degradation governs enhanced sprouting in *HyperDetyr* cells. (A) Representative minimum intensity projections of fluorescent gelatin degraded by endothelial cells after 18 h of invasion, corresponding thresholded cell masks and degraded matrix area (white), and the merge with cell threshold in purple, matrix degradation in green and colocalization in white. Scale bar=8 µm. (B) Quantification shows increased matrix degradation area in the *HyperDetyr* model. (N=8 independent experiments). (C) Maximum displacement is proportional to the area of matrix degradation. (D) Representative maximum intensity projections of angiogenic sprouts in CTRL and *HyperDetyr* models after 8 h and 24 h of invasion, with DMSO or Batimastat treatment. Nuclei were stained with DAPI (yellow) and sprouts with Phalloidin (labeling F-actin, cyan). Scale bar=25 µm. (E, F) Quantification of nuclei per sprout at 8 h (E) and 24 h (F) in the presence of Batimastat shows no significant difference between CTRL and *HyperDetyr* models. Batimastat treatment reduces the number of nuclei per sprout at 24 h in the *HyperDetyr* model (N=14 independent experiments). (G) Representative 3D renders of displacements from 3D TFM experiments around sprouts after 18 h of invasion with DMSO or Batimastat treatment. The bead displacements were quantified (see Methods for details) and the maximum value per sprout was selected. (H) Batimastat slightly reduces maximum displacement in the CTRL model and causes a strong reduction in displacement in the *HyperDetyr* model (n>13, N=3 independent experiments).

We next tested whether enhanced matrix degradation is causally responsible for the increased sprouting observed in *HyperDetyr* cells. HUVECs were treated with the broad-spectrum matrix metalloproteinase (MMP) inhibitors Batimastat or GM6001^53^. In control conditions, *HyperDetyr* cells formed sprouts with similar numbers and nuclear content as CTRL cells at early time points, but significantly greater length **(Fig. 4D, E and Extended Fig. 4A-C)**. Strikingly, upon Batimastat treatment, sprout number and nuclear content remained almost unchanged between CTRL and *HyperDetyr* cells, and importantly, the length advantage of *HyperDetyr* sprouts was fully abolished, restoring values to CTRL levels **(Fig. 4D, E, Extended Fig. 4A-C)**. At later stages (48 h), the elevated cell content per sprout observed in *HyperDetyr* cells was likewise normalized to CTRL upon Batimastat treatment **(Fig. 4D, F)**. Similar results were obtained with the alternative MMP inhibitor GM6001 **(Extended Fig. 4D, E)**. These findings demonstrate that enhanced ECM degradation is required for the detyrosination-dependent sprouting phenotype.

Given the strong correlation between matrix degradation and traction forces, we next asked whether MMP activity also underlies the increased force transmission observed in *HyperDetyr* cells. 3D TFM revealed that the elevated traction forces exerted by *HyperDetyr* sprouts were significantly reduced upon Batimastat treatment **(Fig. 4G, H)**, indicating that ECM degradation is necessary for force generation during invasion. Together, these results identify MMP-dependent matrix remodeling as the principal effector of detyrosination-driven sprouting.

## Discussion

Our study identifies tubulin detyrosination as a positive regulator of endothelial sprouting that promotes multicellular invasion through a coordinated control of migration persistence, ECM remodeling, and force transmission. By engineering endothelial cells with uniform microtubule PTM states, we demonstrate that selective microtubule detyrosination robustly enhances angiogenic sprouting, whereas hyperacetylation does not. Importantly, functional perturbations reveal that this phenotype is driven primarily by increased MMP-dependent ECM degradation, rather than by elevated actomyosin contractility.

Proteolytic ECM remodeling is increasingly recognized as a central determinant of invasive migration across pathological and physiological contexts, including angiogenesis. Our traction force measurements show that, in 3D environments, matrix degradation is not merely permissive but actively supports force transmission. These findings are consistent with previous work demonstrating that localized ECM cleavage remodels matrix architecture and mechanics to facilitate efficient traction force generation and transmission^54–56^. Although crosstalk between microtubules and actomyosin contractility has been implicated in mechanosensitive migration^11,12,57^, our inhibition experiments demonstrate that the enhanced forces observed in detyrosinated cells arise independently of actomyosin activity and instead rely on protease-dependent matrix remodeling. This positions ECM degradation as the dominant effector downstream of microtubule detyrosination during angiogenic invasion.

How microtubule detyrosination promotes enhanced matrix remodeling remains an open question. While intermediate filaments have recently been shown to induce MT1-MMP expression^58^ in invading glioblastoma cells, a plausible mechanism involves altered vesicular trafficking of proteases, leading to increase delivery or retention of MMPs at the cell surface. Microtubule detyrosination is known to modulate interactions with motor machineries implicated in both MMP secretion and endocytosis^15,23,59,60^. The enhanced recruitment of CLASP2 to microtubule plus ends observed in *HyperDetyr* cells may further contribute to this phenotype, as CLASP-mediated microtubule capture at adhesion sites has previously been shown to promote pericellular proteolysis^61^, suggesting a potential mechanistic link between microtubule dynamics and localized ECM degradation. Elucidating how detyrosination couples microtubule behavior to protease trafficking will be an important direction for future work.

Beyond its impact on matrix remodeling, detyrosination fundamentally reshapes microtubule dynamics. Using cellular models in which microtubules are uniformly modified, we show that detyrosination does not globally stabilize the microtubule lattice but instead preserves dynamic instability while selectively promoting persistent growth. Mechanistically, our data support a model in which detyrosination suppresses MCAK-mediated depolymerization events^24^, thereby increasing growth persistence. This dynamic state favors CLASP accumulation at microtubule plus ends, which in turn reinforces persistent growth. Such functional antagonism between MCAK- and CLASP has previously been described in migrating endothelial cells^62^ and may represent a general mechanism sustaining microtubule growth during cell migration. In this context, our findings extend emerging roles for CLASP in mechanically and spatially constrained migration^12,51^, positioning it as a key mediator of persistent and resilient microtubule behavior. While altered microtubule plus-end behavior may influence targeted secretion and matrix degradation, increased microtubule growth persistence may also directly support protrusive activity and directional migration in 3D environments independently of matrix remodeling^52,63–65^. These mechanisms are not mutually exclusive and may act in concert to sustain multicellular sprout extension.

At the cellular level, detyrosination enhances directional migration persistence without affecting migration speed. This contrasts with previously proposed models in which front–rear asymmetry between tyrosinated and detyrosinated microtubules drives persistent migration^27^. Instead, our data reveal that a global enrichment of a dynamic, persistently growing microtubule network is sufficient to stabilize migratory directionality. Such a mechanism is particularly well suited to three-dimensional environments, where sustained protrusive activity and directional persistence are critical for effective invasion.

Our findings also address a gap in endothelial biology regarding the role of tubulin PTMs in angiogenesis. While non-centrosomal microtubules are known to control endothelial polarization during sprouting^7^, the specific contribution of tubulin detyrosination has remained poorly explored. Notably, previous in vivo work identified VASH1-mediated detyrosination as a regulator of venous sprouting and proliferation during lymphovenous development in zebrafish^29^. Whether enrichment of microtubule detyrosination occurs more broadly across vascular beds, potentially promoting sprouting in other contexts, remains to be determined, but our results provide a mechanistic framework linking detyrosination to endothelial invasion through coordinated regulation of microtubule dynamics, migration persistence and matrix remodeling.

In summary, we show that microtubule detyrosination preserves microtubule dynamics while promoting persistent growth, enabling endothelial cells to sustain directional migration and protease-dependent matrix remodeling. By coupling cytoskeletal dynamics to ECM remodeling and force transmission, detyrosinated microtubules endow endothelial cells with enhanced invasive capacity while maintaining responsiveness to angiogenic patterning cues. More broadly, our findings identify the tubulin code as a regulatory layer integrating intracellular mechanics with extracellular remodeling during angiogenesis and suggest that selective modulation of microtubule PTMs may offer new strategies to tune vascular invasion in regenerative and pathological settings.

**Extended Fig. 1.**
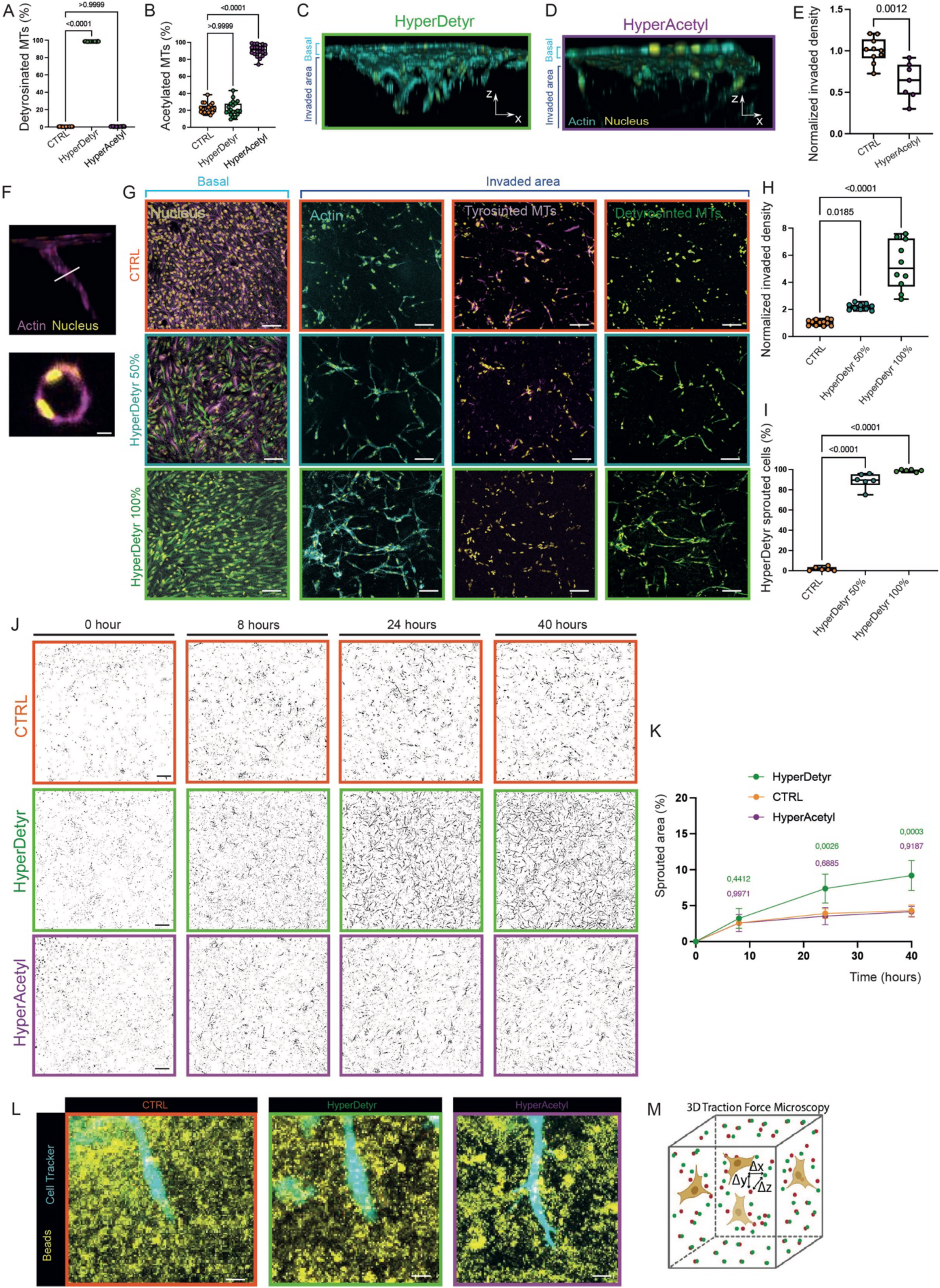
(A,B) Area percentage of acetylated (A) and detyrosinated (B) microtubules normalized to total tubulin in CTRL, *HyperDetyr* and *HyperAcetyl* cells (n=20, N=5 independent experiments). (C,D) 3D volume reconstruction of confocal images showing sprouting HUVECs in *HyperDetyr* (C) and *HyperAcetyl* (D) conditions. Nuclei were stained with DAPI (yellow) and sprouts with Phalloidin (labeling F-actin, cyan). (E) t-test analysis of the normalized invaded density in CTRL and *HyperAcetyl* cells (N>7 independent experiments). The *HyperAcetyl* cells shows a significantly lower normalized invaded density compared to CTRL. (F) 3D confocal reconstruction of a *HyperDetyr* sprout exhibiting a well-defined lumen. Nuclei were stained with DAPI (yellow) and sprouts with Phalloidin (labeling F-actin, magenta). Scale bar=5 µm. (G) Maximum z-projection of the CTRL model (orange), 1:1 co-culture (CTRL: *HyperDetyr*, blue), and *HyperDetyr* model (green) during the sprouting experiment. Scale bar=80 µm. Cells were stained for tyrosinated (magenta) and detyrosinated (green) tubulin, Phalloidin (labeling F-actin, cyan) and nuclei were stained with DAPI (yellow). (H) Quantification of invaded cell density as a function of the proportion of *HyperDetyr* cells used in the sprouting experiment. Invaded cell density increases with higher H*yperDetyr* representation (N=12 independent experiments). (I) In the 1:1 co-culture condition (CTRL/*HyperDetyr*), 85% of invading cells are derived from the *HyperDetyr* population (N=6 independent experiments). (J) Thresholded time-lapse images from live sprouting experiments. Sprouts within the gel were segmented by thresholding, and sprout area was quantified over time. Scale bar=100 µm (K) Quantification of the sprouted area over time reveals a progressive increase in the sprouted area comparing to CTRL model (N=8 independent experiments). (L) Representative maximum intensity projections of fluorescent beads (yellow) overlaid with angiogenic sprouts (cyan) computed around endothelial sprouts after 18 h of invasion. Scale bar=20 µm. (M) Schematic of the 3D traction force microscopy (TFM) experiment. Green beads represent the stressed state and red beads correspond to the relaxed state.

**Extended Fig. 2.**
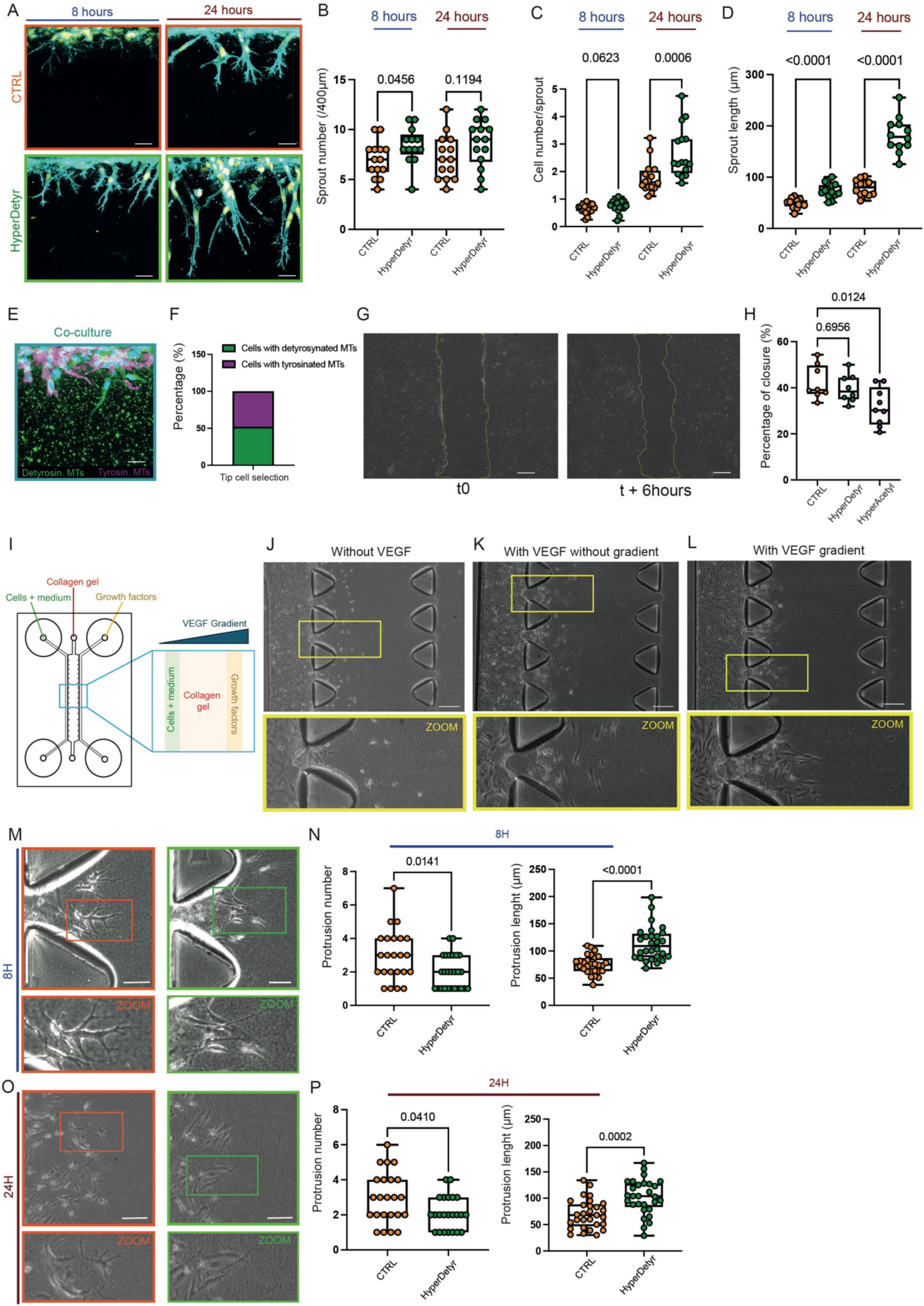
(A) Representative confocal images of sprouting experiments with lateral view at 8 h and 24 h in CTRL and *HyperDetyr* models. Nuclei were stained with DAPI (yellow) and sprouts with Phalloidin (labeling F-actin, cyan). Scale bar=35 µm. (B) Quantification of sprout number indicates no difference between CTRL and *HyperDetyr* conditions at 8 h and 24 h (N=14 independent experiments). (C) The number of cells per sprout significantly increases in *HyperDetyr* cells after 24 h of sprouting (N=14 independent experiments). (D) Sprout length is consistently increased in the *HyperDetyr* model at both time points (N=14 independent experiments). (E) Confocal image of a co-culture sprouting assay after 6 h of sprouting. Cells were stained for tyrosinated (magenta) and detyrosinated (green) tubulin and nuclei were stained with DAPI (blue). Scale bar=20 µm. (F) The percentage of tip cells enriched in tyrosinated or detyrosinated microtubules was quantified. No preferential tip cell selection is observed between tyrosinated microtubule-enriched CTRL cells (purple) and detyrosinated microtubule-enriched *HyperDetyr* cells (green) (N=3 independent experiments). (G) Brightfield images of a 2D scratch wound assay at 0 h and after 6 h of migration. Scale bar = 300 µm. (H) Quantification of wound closure after 6 h shows no significant change for the *HyperDetyr* model, whereas closure is reduced in the *HyperAcetyl* condition (N>7 independent experiments). (I) Schematic of the microsystem used to generate a VEGF gradient. The system consists of three parallel channels: a central channel filled with collagen gel, flanked by two lateral channels, one containing control medium and the other supplemented with growth factors, thereby generating a gradient across the gel. (J,K,L) Brightfield images of invasion assays under different VEGF conditions: without VEGF (J), with no gradient but a uniform VEGF stimulation (K) and with a VEGF gradient (L). Yellow rectangles indicate the areas selected for higher-magnification views shown above. Cells show increased invasion in response to VEGF (K,L) and orient their migration toward the gradient (L); Scale bar=200 µm. (M,O) Brightfield images of CTRL and *HyperDetyr* models after 8 h (M) and 24 h (O) in 3D collagen. Scale bar=60 µm. (N,P) Quantification of protrusive activity. The number of protrusions is reduced in *HyperDetyr* cells at both 8 h (N) and 24 h (P), while protrusion length is increased in the *HyperDetyr* model at both time points (n>20, N=5 independent experiments). Orange (CTRL) and green (*HyperDetyr*) rectangles indicate the areas selected for higher-magnification views shown above.

**Extended Fig. 3.**
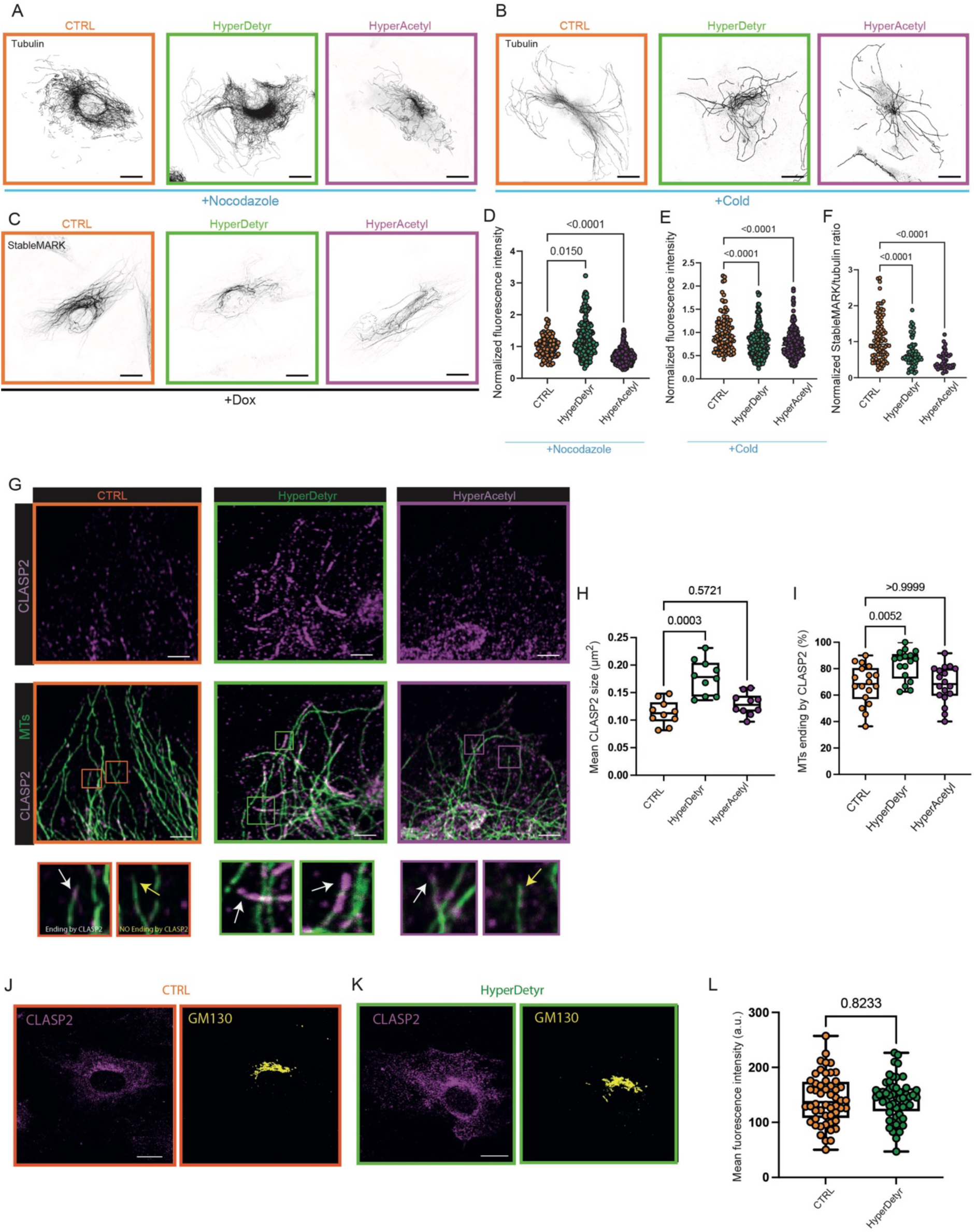
(A-C) Representative immunofluorescence images of CTRL, *HyperDetyr* and *HyperAcetyl* models following (A) nocodazole treatment, (B) cold-induced depolymerization, and (C) StableMARK live staining. Cells were stained for α-tubulin. Scale bar=5 µm. (D) Quantification of normalized fluorescence intensity after nocodazole treatment reveals increased intensity in the *HyperDetyr* model and reduced intensity in the *HyperAcetyl* model relative to CTRL (n>97, N=3 independent experiments). (E) Quantification of normalized fluorescence intensity after cold-induced depolymerization shows decreased intensity in both *HyperDetyr* and *HyperAcetyl* models compared to CTRL (n>99, N=3 independent experiments). (F) The ratio of total tubulin to StableMARK-labeled tubulin is likewise reduced in *HyperDetyr* and *HyperAcetyl* models relative to CTRL (n>65, N=3 independent experiments). (G) Confocal images of CLASP2 immunostaining in CTRL, *HyperDetyr* and *HyperAcetyl* cells. Cells were stained for α-tubulin (green) and CLASP2 (magenta). Scale bar=2 µm. White arrows indicate microtubules with CLASP2-positive ends, whereas yellow arrows indicate microtubules without detectable CLASP2 signal at their ends. Quantification of CLASP2 signal: (H) average CLASP2 size at the end of microtubules and (I) percentage of microtubules ending with CLASP2. All parameters increase in the *HyperDetyr* model compared to CTRL and *HyperAcetyl* conditions (N>10 independent experiments). (J,K) Confocal images of CLASP2 immunostaining (magenta) in CTRL (J) and *HyperDetyr* (K) models, showing Golgi-associated signal (GM130, yellow). Scale bar=5 µm. (L) Quantification of CLASP2 mean fluorescence intensity at the Golgi reveals no significant difference between CTRL and *HyperDetyr* conditions (n>50, N=3 independent experiments).

**Extended Fig. 4.**
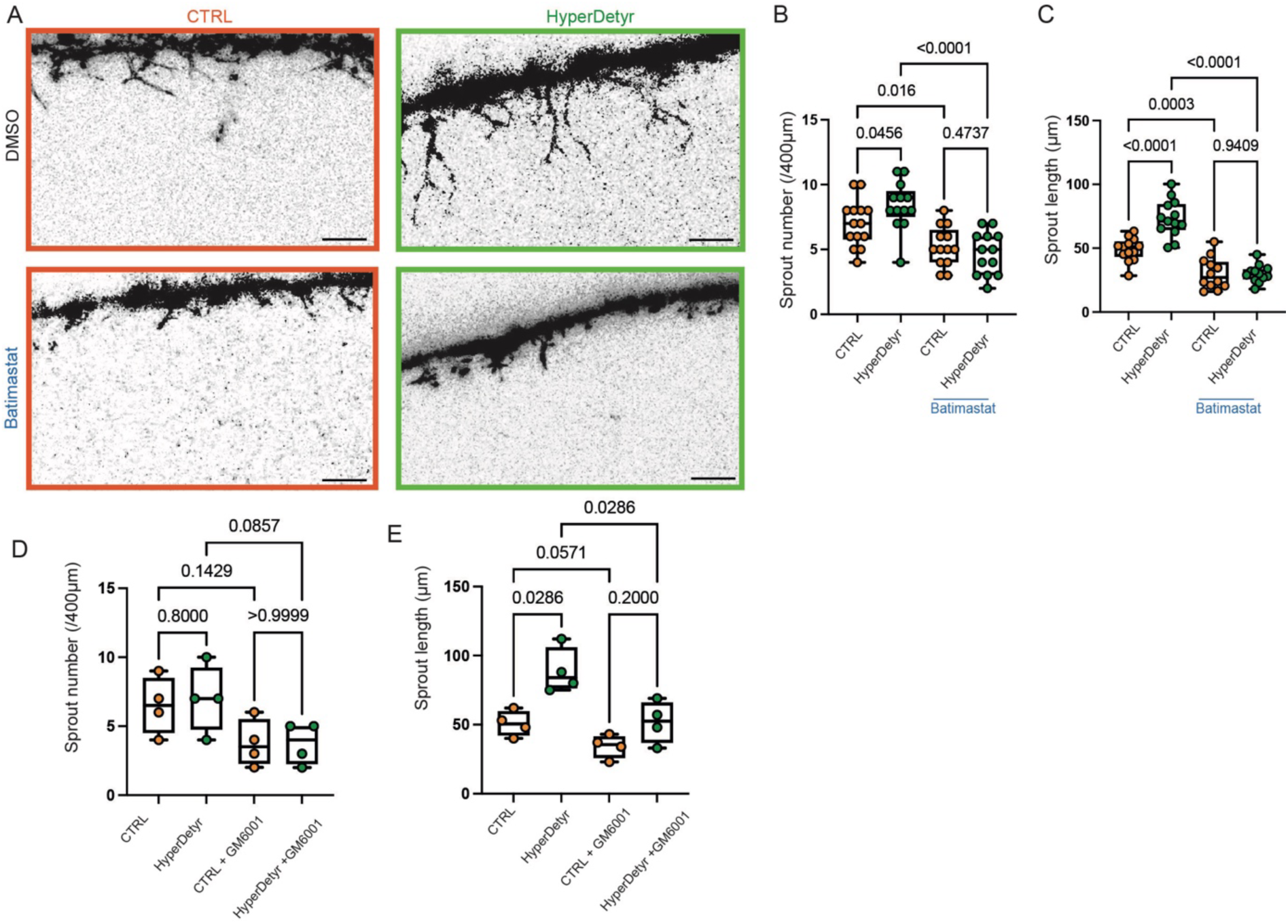
(A) Representative maximum intensity projections of angiogenic sprouts in CTRL and *HyperDetyr* models with DMSO or Batimastat treatment. Sprouts were stained with Phalloidin (labeling F-actin). Scale bar=50 µm. (B,C) Batimastat treatment slightly reduces (B) sprout number and (C) length in the CTRL model, but significantly impairs sprouting in the *HyperDetyr* model, reducing both metrics to levels comparable with CTRL (n>13, N=6 independent experiments). (D) Sprout number and (B) sprout length are reduced in both CTRL and *HyperDetyr* models in the presence of GM6001, another broad-spectrum MMP inhibitor, confirming the requirement for MMP activity in sprouting (N=4 independent experiments).

## Materials & methods

### Plasmid construction

All constructs were generated using the In-Fusion Cloning® HD kit (Takara, cat. no. 638948) to insert the gene of interest into the pLVX-IRES-Puro plasmid (Clontech). For StableMARK, the pLVX-TetOne-Puro plasmid (Clontech) was used for inducible expression. Plasmids were previously digested with EcoRI-HF and BamHI-HF (New England Biolabs, cat. nos. R3101L and R3136L).

The coding sequences were amplified from HUVEC cDNA, and the StableMARK sequence was amplified from the eGFP-KIF5B-Rigor plasmid (Addgene plasmid #172204). DNA fragments were purified using the Monarch® DNA Gel Extraction Kit (New England Biolabs, cat. no. T1020L) according to the manufacturer’s instructions.

### Cell culture

Human umbilical vein endothelial cells (HUVECs; PromoCell, cat. no. C-12203) were cultured in Endothelial Cell Growth Medium (PromoCell, cat. no. C-22010) at 37 °C and 5% CO₂. Human embryonic kidney cells (HEK293T) were cultured in DMEM/F12 (Capricorn, cat. no. DMEM-12-A) supplemented with 10% fetal bovine serum (Biowest, cat. no. S1810) and 1% penicillin–streptomycin (Sigma, cat. no. P4333) at 37 °C and 5% CO₂.

### Lentivirus production

For lentiviral particle production, HEK293T cells were transfected in 10-cm dishes at 80% confluence. The transfection mix consisted of 15 µg of the plasmid containing the construct of interest, 10 µg of psPAX2 (plasmid expressing HIV-1 gag and pol polyproteins), and 5 µg of pMD2.G (plasmid expressing VSV-G envelope proteins) diluted in 1.2 mL of Opti-MEM I medium. The mixture was gently vortexed and incubated for 5 min at room temperature. MaxPEI (Polysciences, cat. no. 24765) was added at a 3:1 ratio relative to the amount of DNA and incubated for 15 min at room temperature before being added dropwise to the cells. The medium was refreshed the following day.

Viral supernatants were harvested on days 3 and 4, filtered through a 0.45 µm syringe filter, and precipitated using a 5x PEG solution (100 g PEG 6000 and 6 g NaCl dissolved in ddH₂O to a final volume of 250 mL, pH 7.2). The mixture was gently mixed, incubated overnight at 4°C, and centrifuged for 30 min at 1,500 × g at 4°C. The supernatant was discarded, and the viral pellet was resuspended in 200 µL of sterile dPBS (Capricorn, cat. no. CA PBS-1A). Viral particle suspensions were aliquoted and stored at -80°C.

### Lentiviral transduction

We generated HUVEC cell lines overexpressing VASH1-SVBP (HyperDetyr model), αTAT1 (HyperAcetyl model), Tet-On GFP (control model), EB3-mScarlet, or Tet-On StableMARK-mNeonGreen, using transduction. Polyclonal populations were used for all experiments, with independent transduction performed for replicates. Cells were seeded in 6-well plates. The medium was replaced 1 h prior to infection with fresh medium supplemented with polybrene (stock: 10 mg/mL; diluted 1:1,250 in culture medium), and 2-5 µL of viral suspension was added to the cells. Forty-eight hours after infection, the medium was replaced with medium containing 1.5 µg/mL puromycin, and selection was maintained for 2-3 days.

### BrdU cell proliferation assay

The bromodeoxyuridine (BrdU) Cell Proliferation ELISA Kit (Abcam, cat. no. ab126572) was used to quantify DNA synthesis and cell proliferation. BrdU was added to compound-treated and control wells to allow incorporation into the DNA of proliferating cells. After 48 h of culture (with BrdU added during the last 24 h), cells were fixed, washed, and processed according to the manufacturer’s ELISA protocol. BrdU incorporation in control cells was set to 100%.

### 3D collagen invasion assay

The collagen matrix consisted of a 2.5 mg/mL collagen solution diluted in Endothelial Cell Growth Medium supplemented with 10% fetal bovine serum (FBS), 50 ng/mL VEGF-165 (vascular endothelial growth factor; PeproTech, cat. no. 100-20-50UG), 50 ng/mL bFGF (basic fibroblast growth factor; PeproTech, cat. no. 100-18C), and 50 ng/mL PMA (phorbol 12-myristate 13-acetate; Sigma-Aldrich, cat. no. P8139-5MG). The solution was buffered with 1/10 Hank’s Balanced Salt Solution (HBSS; Sigma-Aldrich, cat. no. 55021C), 1/40 HEPES, and NaOH to adjust the pH to 7.0-7.5.

The collagen mixture was either placed in a pre-warmed Lab-Tek™ culture chamber (Thermo Fisher Scientific, cat. no. 154534PK) or pipetted into a pre-cut imaging chamber (Invitrogen Secure-Seal™, cat. no. S24732) adhered to the glass bottom of a pre-warmed 24-well plate (Cellvis, cat. no. P24-0-N), and allowed to polymerize at 37 °C for 10 min. For imaging chambers, 40,000 HUVECs per chamber were seeded to obtain a confluent monolayer by incubating the plate vertically for 1 h at 37 °C and 5% CO₂, allowing cell attachment onto the collagen gel surface. Plates were then placed horizontally and incubated for the indicated time in complete growth medium. For Lab-Tek chambers, 90,000 HUVECs were seeded per well.

To inhibit matrix metalloproteinase (MMP) activity, cells were treated throughout the experiments with 5 µM Batimastat (Sigma-Aldrich, cat. no. 196440) or 15 µM GM6001 (Sigma-Aldrich, cat. no. CC10).

Rho-associated kinase (ROCK) activity was inhibited using Y-27632 (Sigma-Aldrich, cat. no. Y0503) at a final concentration of 10 µM.

### VEGF gradient assay

The idenTx 3 microfluidic chips were purchased from AIM Biotech. The chip consists of three channels. The gel channel of the microfluidic chips was filled with collagen type I hydrogel. After 5 minutes of polymerization, 10 μL of an endothelial cell suspension (3 × 10⁶ cells/mL) was introduced into one of the lateral media channels. The chips were then incubated in a humidified chamber to allow cell adhesion to the collagen interface. Subsequently, both media channels were filled with 120 μL of EGM-2 medium, either supplemented or not with angiogenic growth factors. This configuration established a chemotactic gradient across the gel, with the left channel without growth factors and the right channel providing the pro-angiogenic stimulus.

### Monolayer wound healing assay

A confluent HUVEC monolayer was scratched using a sterile P200 tip to create a cell-free zone. mages were acquired immediately after scratching and after 8 h. Cell migration was quantified by calculating the percentage of wound closure using ImageJ.

### Stamp fabrication and microcontact printing for 1D migration

Microstripes of 2 µm were produced using a silicon master fabricated by deep reactive-ion etching from a chromium photomask (Toppan Photomask). The silicon surface was passivated with a fluorosilane (tridecafluoro-1,1,2,2-tetrahydrooctyl-1-trichlorosilane; Gelest) for 30 min. Polydimethylsiloxane (PDMS) microstamps were cast from the silanized master using Sylgard 184 (Dow Corning) and cured for 4 h at 60 °C.

PDMS stamps were oxidized in a UV/O₃ oven for 7 min and incubated for 1 h at room temperature with a fibronectin solution (18 µg/mL, human plasma fibronectin; Sigma-Aldrich). After drying under filtered air, stamps were gently applied to flat PDMS-coated glass coverslips. Non-printed areas were passivated by incubating the substrates for 5 min in 1% Pluronic F-127 (BASF), followed by three washes with PBS.

Time-lapse microscopy was used to monitor cell migration on 1D micropatterned lines. Cells were tracked using the MATLAB plugin CellTracker^66^ and quantitative analyses were performed with GraphPad Prism. To enable live-cell imaging over at least 8 h, cell nuclei were labeled with Hoechst dye.

### 3D traction force microscopy image acquisition

Confocal images were acquired at a resolution of 512 x 512 pixels using a 20x/0.4 NA dry objective with 2x zoom, a 56.6 µm pinhole corresponding to 1 Airy unit, and bidirectional scanning with phase correction. Excitation wavelengths were 488 nm and 638 nm. Emission detection windows for photomultiplier tubes were 498-538 nm and 673-704 nm, respectively. The voxel size was set to 0.568 x 0.568 x 1 µm.

All traction force microscopy (TFM) experiments were performed on a Leica Stellaris 8 confocal microscope equipped with a temperature-controlled stage set to 37 °C and a humidified 5% CO₂ atmosphere. Regions of interest were first identified and registered. Imaging was initiated by acquiring the stressed (force-exerting) state. Cells were then treated with 4 µM cytochalasin D (dissolved in DMSO; Sigma-Aldrich) for 1 h, after which the relaxed (force-free) state was acquired at the same positions.

### 3D traction force microscopy analysis

Three-dimensional TFM datasets were processed using the TFMLAB software toolbox^67^. Bead and sprout images were filtered and enhanced, and sprout images were binarized to generate a 3D cell mask. Stage drift between stressed and relaxed acquisitions was corrected by phase correlation. Three-dimensional bead displacements were computed using free-form deformation-based image registration. Regions of interest around sprout tips were subsequently selected for analysis.

### Matrix degradation assay

Gelatin labeled with Alexa Fluor 647 (Thermo Fisher Scientific) was incorporated at twice the final concentration into the collagen solution to generate a uniformly fluorescent hydrogel. Following angiogenic invasion, hydrogels were imaged using a 647-nm laser to visualize degraded regions, which appeared as dark areas. Images were processed using a disk-shaped top-hat filter to correct background inhomogeneity, followed by Gaussian smoothing and contrast stretching. A minimum-intensity projection was generated to obtain the complementary image. Degraded regions were binarized using a manually adjusted intensity threshold, and the degradation area was quantified around the sprout of interest and normalized to the number of nuclei.

### Immunostaining 2D

Cells were seeded on glass coverslips placed in 24-well plates. Fixation was performed using either 100% cold methanol for 10 min at -20 °C or 4% paraformaldehyde (PFA) for 15 min at room temperature. Cells were washed twice with PBS and permeabilized with PBS containing 0.15% Triton X-100 for 3 min at room temperature with agitation. After two additional PBS washes, cells were blocked with PBS containing 0.05% Tween-20 and 2% bovine serum albumin (BSA; Roth, cat. no. 8076.4) for 1 h.

Coverslips were removed from the wells and placed on parafilm in a humid chamber. Samples were incubated for 1.5 h with 40 µL of primary antibodies diluted in blocking buffer, washed four times for 5 min with PBS/0.05% Tween-20, and incubated for 1 h with 40 µL of secondary antibodies. After washing four times with PBS/0.05% Tween-20 and once with PBS followed by water, coverslips were dried and mounted using Mowiol mounting medium containing DAPI.

For microtubule depolymerization assays, cells were treated with 500 nM nocodazole (Sigma-Aldrich, cat. no. M1404) for 2 h at room temperature or incubated at -20 °C for 5 min. Cytosolic contents were pre-extracted prior to fixation to remove soluble tubulin by incubating cells for 1 min at 37 °C in extraction buffer (80 mM PIPES, pH 6.8; 1 mM MgCl₂; 4 mM EGTA; 0.3% Triton X-100; 0.1% glutaraldehyde).

### 3D

Cells embedded in 3D matrices were fixed with 4% PFA for 20 min at room temperature and washed twice with PBS:glycine. Permeabilization was performed for 45 min at room temperature with gentle agitation (50 rpm) using PBS:glycine containing 0.5% Triton X-100. Samples were blocked for 1 h at room temperature with agitation in IF wash solution (PBS containing 7 mM NaN₃, 0.2% Triton X-100, and 0.05% Tween-20) supplemented with 1% BSA.

Phalloidin-iFluor 647™ (Abcam, cat. no. ab176759) and DAPI were diluted in blocking solution and incubated with the gels for 2 h. Gels were washed four times for 20 min with IF wash solution under agitation, followed by two washes with water. Samples were briefly air-dried at room temperature and mounted using fluorescence mounting medium (Dako, cat. no. 302380-02).

**Table 1:** antibodies.

| Antibodies | Source | Reference |
| --- | --- | --- |
| Anti-alpha tubulin, rabbit monoclonal | Abcam | ab52866 |
| Anti-tyrosinated alpha tubulin (YL1/2), rat monoclonal | Thermofisher | MA1-80017 |
| Anti-detyrosinated alpha tubulin, rabbit polyclonal | Abcam | ab48389 |
| Anti-acetylated alpha tubulin, mouse monoclonal | Sigma-Aldrich | T6793 |
| Anti-EB1, mouse monoclonal | BD Transduction Laboratories | 610535 |
| Anti-CLASP2, rat monoclonal | Abcam | ab95373 |
| Anti-GM130, mouse monoclonal | BD Transduction Laboratories | 610823 |
| Phalloidin-iFluor 647 | Abcam | ab176759 |
| Alexa Fluor Plus 647 donkey anti-rat | Invitrogen | A48272 |
| Alexa Fluor 546 donkey anti-mouse | Invitrogen | A10036 |
| Alexa Fluor 488 goat anti-rabbit | Invitrogen | A11008 |
| Alexa Fluor 488 goat anti-mouse | Invitrogen | A11001 |
| Alexa Fluor 594 goat anti-mouse | Invitrogen | A11005 |

### Microscopy

Fluorescence microscopy images were taken using a Leica Stellaris 5 confocal microscope equipped with HC PL APO CS2 objectives (63x/1.4 (oil) or 20x/0.75 (air)), or a Leica Stellaris 8 confocal microscope equipped with HC PL APO 20x/0.75 (oil) objective. Both were equipped with an OKO lab live cell culture incubator to main the temperature at 37 °C with 5% CO_2_. For imaging of EB dynamics, an 8Khz resonant scanner on the Leica Stellaris 5 was used together with a Dynamic Signal Enhancement of 8, allowing acquisition every 50 ms.

### Image preparation and analysis

For image processing, ImageJ was used for level and contrast adjustments, maximum-intensity projections and thresholding. The resulting binary masks were then employed for particle analysis and detection.

Quantification of microtubule post-translational modifications was performed either by measuring the fluorescence intensity of each specific modification normalized to total tubulin fluorescence, or by calculating the surface area of thresholded post-translational modifications normalized to the total tubulin area.

Kymographs of microtubule plus-end dynamics were generated using the KymoResliceWide plugin in ImageJ and analyzed using the same software. Only growth events that both initiated and terminated within the acquisition period were included in the analysis. Growth velocity was calculated for each individual event and subsequently averaged. Microtubule persistence length was defined as the distance traveled during a growth phase, from initiation to catastrophe.

EB3 comets and CLASP2 stretches were automatically detected on thresholded images using the Particle Analysis plugin in ImageJ. EB3 comet and CLASP2 stretch were quantified and normalized to the total cell surface area.

All mentioned ImageJ plugins have source code available and are licensed under open-source license.

### Statistical analysis

GraphPad Prism was used to create all graphs and to perform statistical analyses. Kruskal-Wallis test with Dunn’s multiple comparisons test were performed when comparing 3 samples after outliers were removed using ROUT method (Q=1%). Mann-Whitney test was performed when comparing 2 samples. For the quantification of the sprouted area over time (Extended Fig. 1K) a two-way ANOVA test was performed.

